# XBB.1.5 monovalent mRNA vaccine booster elicits robust neutralizing antibodies against emerging SARS-CoV-2 variants

**DOI:** 10.1101/2023.11.26.568730

**Authors:** Qian Wang, Yicheng Guo, Anthony Bowen, Ian A. Mellis, Riccardo Valdez, Carmen Gherasim, Aubree Gordon, Lihong Liu, David D. Ho

**Affiliations:** Aaron Diamond AIDS Research Center, Columbia University Vagelos College of Physicians and Surgeons, New York, NY, USA; Division of Infectious Diseases, Department of Medicine, Columbia University Vagelos College of Physicians and Surgeons, New York, NY, USA; Department of Pathology and Cell Biology, Columbia University Vagelos College of Physicians and Surgeons, New York, NY, USA; Department of Pathology, University of Michigan, Ann Arbor, MI, USA; Department of Epidemiology, University of Michigan, Ann Arbor, MI, USA; Department of Microbiology and Immunology, Columbia University Vagelos College of Physicians and Surgeons, New York, NY, USA

**Keywords:** COVID-19, SARS-CoV-2, Omicron subvariants, XBB.1.5 monovalent mRNA vaccine, HV.1, HK.3, JD.1.1, JN.1, Immunological imprinting

## Abstract

COVID-19 vaccines have recently been updated with the spike protein of SARS-CoV-2 XBB.1.5 subvariant alone, but their immunogenicity in humans has yet to be fully evaluated and reported, particularly against emergent viruses that are rapidly expanding. We now report that administration of an updated monovalent mRNA vaccine (XBB.1.5 MV) to uninfected individuals boosted serum virus-neutralization antibodies significantly against not only XBB.1.5 (27.0-fold) and the currently dominant EG.5.1 (27.6-fold) but also key emergent viruses like HV.1, HK.3, JD.1.1, and JN.1 (13.3-to-27.4-fold). In individuals previously infected by an Omicron subvariant, serum neutralizing titers were boosted to highest levels (1,504-to-22,978) against all viral variants tested. While immunological imprinting was still evident with the updated vaccines, it was not nearly as severe as the previously authorized bivalent BA.5 vaccine. Our findings strongly support the official recommendation to widely apply the updated COVID-19 vaccines to further protect the public.

## Introduction

Although the World Health Organization (WHO) has announced the conclusion of the emergency phase of the COVID-19 pandemic (WHO, 2023), SARS-CoV-2 continues to spread and evolve (Wang et al., 2023d; Wang et al., 2023f). Emerging viral variants increasingly evade host immunity acquired through vaccination, natural infection, or both, thereby posing a persistent threat to public health (Carabelli et al., 2023). In particular, the emergence of Omicron XBB subvariants has dramatically reduced the efficacy of both SARS-CoV-2 wildtype monovalent and bivalent (wildtype + Omicron BA.5) mRNA vaccines (Wang et al., 2023g), prompting the United States Food and Drug Administration (FDA) to authorize monovalent XBB.1.5-spike-based vaccines for individuals who are older than 6 months, starting in the Fall of 2023 (FDA, 2023). Preliminary studies indicate that the updated monovalent vaccines substantially boosted serum virus-neutralizing antibody titers against previously dominant Omicron subvariants, such as XBB.1.5 and EG.5.1 (Chalkias et al., 2023; Kosugi et al., 2023; Modjarrad et al., 2023; Patel et al., 2023; Stankov et al., 2023), but their impact on viral variants that have subsequently emerged remains to be determined.

A number of SARS-CoV-2 Omicron subvariants have emerged recently, with several gaining traction in different parts of the globe (Elbe and Buckland-Merrett, 2017) (**Figure 1A**). Notably, over 40% of the new infections in Asia are attributed to HK.3, whereas HV.1 constitutes upwards of 34% of the new cases in North America. In Europe, subvariants JD.1.1, BA.2.86, and JN.1 are expanding, each accounting for 6.5%, 4.1%, and 11.8%, respectively. HV.1, HK.3, and JD.1.1 have evolved from the XBB lineage, while JN.1 is a slight variant of BA.2.86 (Wang et al., 2023d), which emerged independently from Omicron BA.2 (**Figure 1B**). Genetically, these subvariants have accumulated additional mutations in their spike proteins. Compared to the recently dominant EG.5.1, HK.3 possesses a unique mutation, L455F, while HV.1 carries two more mutations, F157L and L452R (**Figure 1C**). JD.1.1 has three spike substitutions on top of those found in XBB.1.5, including the so-called “flip mutations” L455F and F456L as well as A475V. Moreover, JN.1 has an additional L455S mutation on the spike protein of BA.2.86 (**Figure 1D**). Interestingly, the aforementioned mutations reside predominantly in the class-1 epitope cluster (Barnes et al., 2020) on the receptor-binding domain (RBD) of spike **(Figure S1)**. In this study, we examined the clinical outcome of an XBB.1.5 monovalent mRNA vaccine boost on serum neutralizing antibodies against these emerging and expanding SARS-CoV-2 Omicron subvariants.

**Figure 1.**
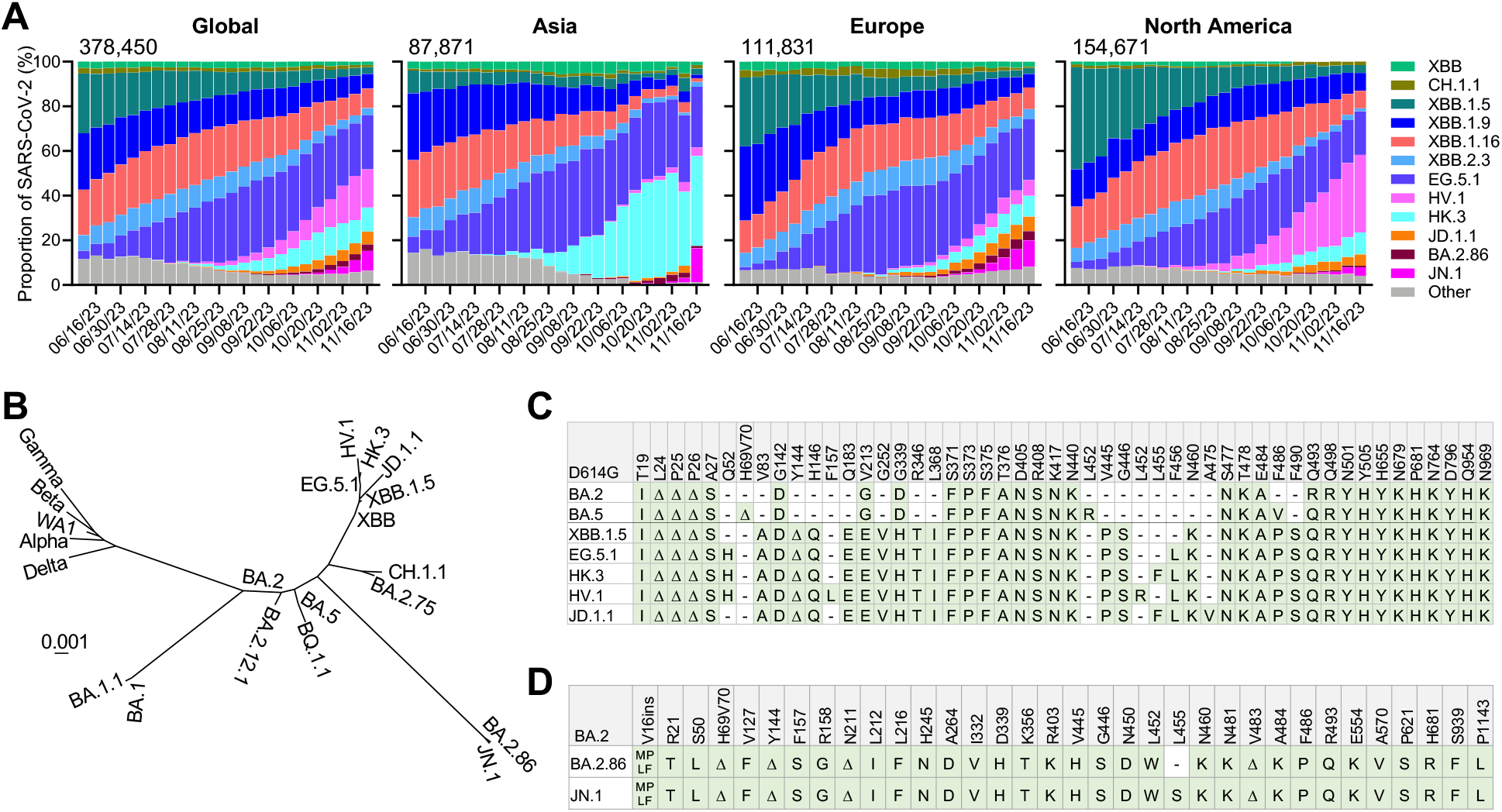
Emergence of novel SARS-CoV-2 variants. **A**. Frequencies of SAR-CoV-2 Omicron subvariants in the denoted time period. Data were obtained from the Global Initiative on Sharing All Influenza Data (GISAID) (Elbe and Buckland-Merrett, 2017). The values in the upper left corner of each box denote the cumulative number of SAR-CoV-2 sequences deposited. **B**. Phylogenetic tree based on spike proteins of SARS-CoV-2 variants. **C**. Spike protein mutations in BA.2, BA.5, XBB.1.5, EG.5.1, HK.3, HV.1, and JD.1.1 relative to D614G. **D**. Spike protein mutations in BA.2.86 and JN.1 relative to BA.2. See also **Figure S1**.

## Results

### Serum neutralization of emerging viral subvariants after an XBB.1.5 mRNA booster

To investigate the neutralizing antibody responses induced by XBB.1.5 mRNA monovalent vaccines against currently circulating and newly emerged subvariants, serum samples from 60 individuals across three different cohorts were collected. To accurately represent real-world conditions, all participants had previously received three to four doses of wildtype monovalent mRNA vaccines followed by one dose of a BA.5 bivalent mRNA vaccine. The three cohorts were 1) individuals with no recorded SARS-CoV-2 infections who received an XBB.1.5 monovalent vaccine booster (“XBB.1.5 MV”); 2) individuals with a recent XBB infection who did not receive an XBB.1.5 vaccine booster (“XBB infx”); and 3) individuals with a prior Omicron infection who also received an XBB.1.5 monovalent vaccine booster (“Omicron infx + XBB.1.5 MV”). The final cohort was further divided into two subgroups: subgroup 1 with a documented infection prior to 2023 (pre-XBB Omicron infection), and subgroup 2 with a documented infection after February 2023 (XBB infection). Detailed demographics of study participants and their vaccination and infection histories are summarized in **Tables S1 and S2. Figure 2A** depicts the timeline of vaccine administration, SARS-CoV-2 infection, and serum collection for each cohort, and the time intervals between serum samples pre and post XBB.1.5 infection or monovalent vaccine boost are similar.

**Figure 2.**
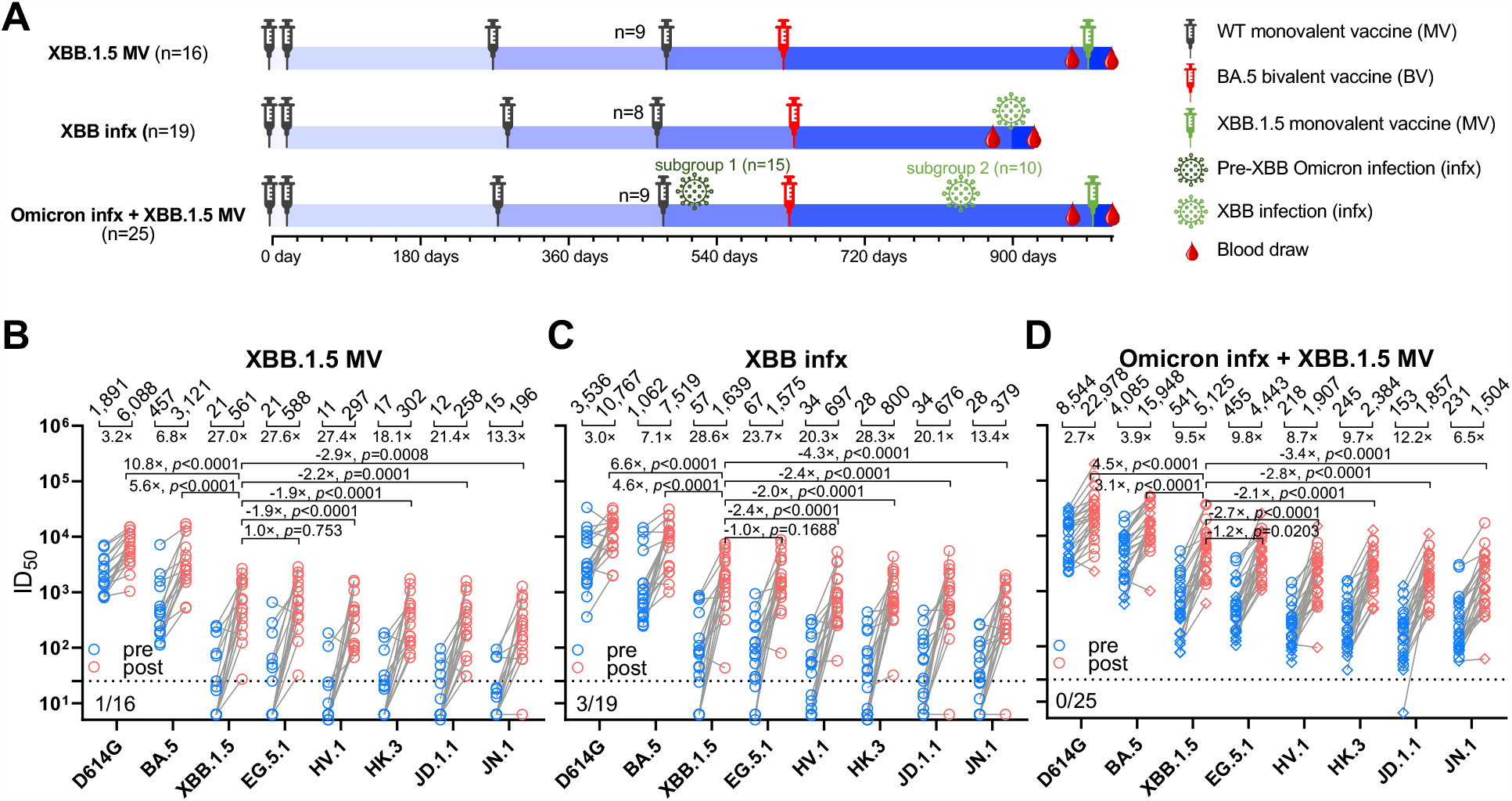
Neutralizing antibody titers before and after an XBB.1.5 mRNA booster, XBB infection, or both. **A**. Timeline representation of vaccine administration, SARS-CoV-2 infection, and serum collection intervals for each clinical cohort. Indicated timepoints represent the median in days for each cohort, with day 0 defined as the day of the initial SARS-CoV-2 vaccination. Numbers of participants for each group receiving a fourth wildtype (WT) monovalent vaccine (MV) is indicated. Other vaccine doses were received by all participants in each cohort. 15 participants from the “Omicron infx + XBB.1.5 MV” cohort had a pre-XBB Omicron infection (subgroup 1), while the other 10 had XBB infection (subgroup 2). n, sample size. **B-D**. Serum virus-neutralizing titers (ID_50_) of the cohorts against the indicated SARS-CoV-2 pseudoviruses. Geometric mean ID_50_ titers (GMT) are shown along with the fold-change between pre and post (MV or infx) serum samples. Horizontal bars show the fold change in GMT following XBB MV or infection between XBB.1.5 and all other viruses tested. The dotted line represents the assay limit of detection (LOD) of 25. Numbers under the dotted lines are non-responders to XBB MV or infection (<3-fold increase in ID_50_ titers between pre- and post-XBB sera across all the viruses tested). In the “Omicron infx + XBB.1.5 MV” cohort, subgroups 1 and 2 are shown in rhombuses and circles, respectively. Statistical analyses were performed by Wilcoxon matched-pairs signed-rank tests. See also **Table S1 and S2 and Figure S2 and S3**.

VSV-pseudotyped viruses were constructed for the emerging subvariants HV.1, HK.3, JD.1.1, and JN.1 as well as D614G, BA.5, XBB.1.5, EG.5.1. These pseudoviruses were then subjected to neutralization assays by pre and post serum samples from the cohorts. In the “XBB.1.5 MV” cohort, the post-vaccination sera showed a 3.2-fold increase in neutralizing ID_50_ (50% inhibitory dilution) titers against D614G and a 6.8-fold increase against BA.5, compared to pre-vaccination sera (**Figure 2B**). A larger increase in ID_50_ titers was observed between pre and post sera against XBB.1.5, EG.5.1, HV.1, HK.3, JD.1.1, and JN.1, ranging from 13.3 to 27.6-fold. The magnitude of these boosts was similar to those found for the “XBB infx” cohort (**Figure 2C**), which exhibited a 3.0-fold increase against D614G, a 7.1-fold increase against BA.5, and 13.4-to-28.6-fold increases against XBB.1.5 and subsequent Omicron subvariants. Not surprisingly, sera from the “Omicron infx + XBB.1.5 MV” cohort displayed highest neutralization titers overall but smaller increases (**Figure 2D**), largely attributable to higher titers in pre-vaccination samples due to a prior Omicron infection. Notably, the increase in neutralization activity following the XBB.1.5 monovalent vaccine booster was again much more pronounced against XBB.1.5 and newer Omicron subvariants (6.5-to-12.2-fold), compared to D614G and BA.5 (2.7-fold and 3.9-fold, respectively). No significant differences in neutralizing titers after vaccination were observed between subgroups 1 and 2 in the last cohort (**Figure 2D and Figure S2**).

After XBB.1.5 vaccination or infection across all three cohorts, the serum neutralization ID_50_ titers against D614G were the highest, ranging from 6,088 to 22,978, followed by those against BA.5, ranging from 3,121 to 15,948 (**Figures 2B, 2C, and 2D**). Compared to BA.5, XBB.1.5 was significantly more (3.1-to-5.6-fold) resistant to neutralization by these sera, whereas it was minimally more (1.0-to-1.2-fold) sensitive than EG.5.1. Serum neutralization titers against newly emerged subvariants HV.1, HK.3, and JD.1.1 were quite similar, but significantly lower than that against XBB.1.5 by 1.9-to-2.8-fold. Overall, serum titers against JN.1 were the lowest, by 2.9-to-4.3-fold relative to titers against XBB.1.5, which is expected given the exposure histories of these cohorts. Importantly, the absolute neutralization titers were robust against all viral variants tested for serum samples after XBB.1.5 vaccination or infection (**Figures 2B, 2C, and 2D**), and the potency and breadth of the antibody boosts were similar for the two XBB.1.5 monovalent mRNA vaccines from different manufacturers, Moderna and Pfizer (**Figures 3A and 3B**).

**Figure 3.**
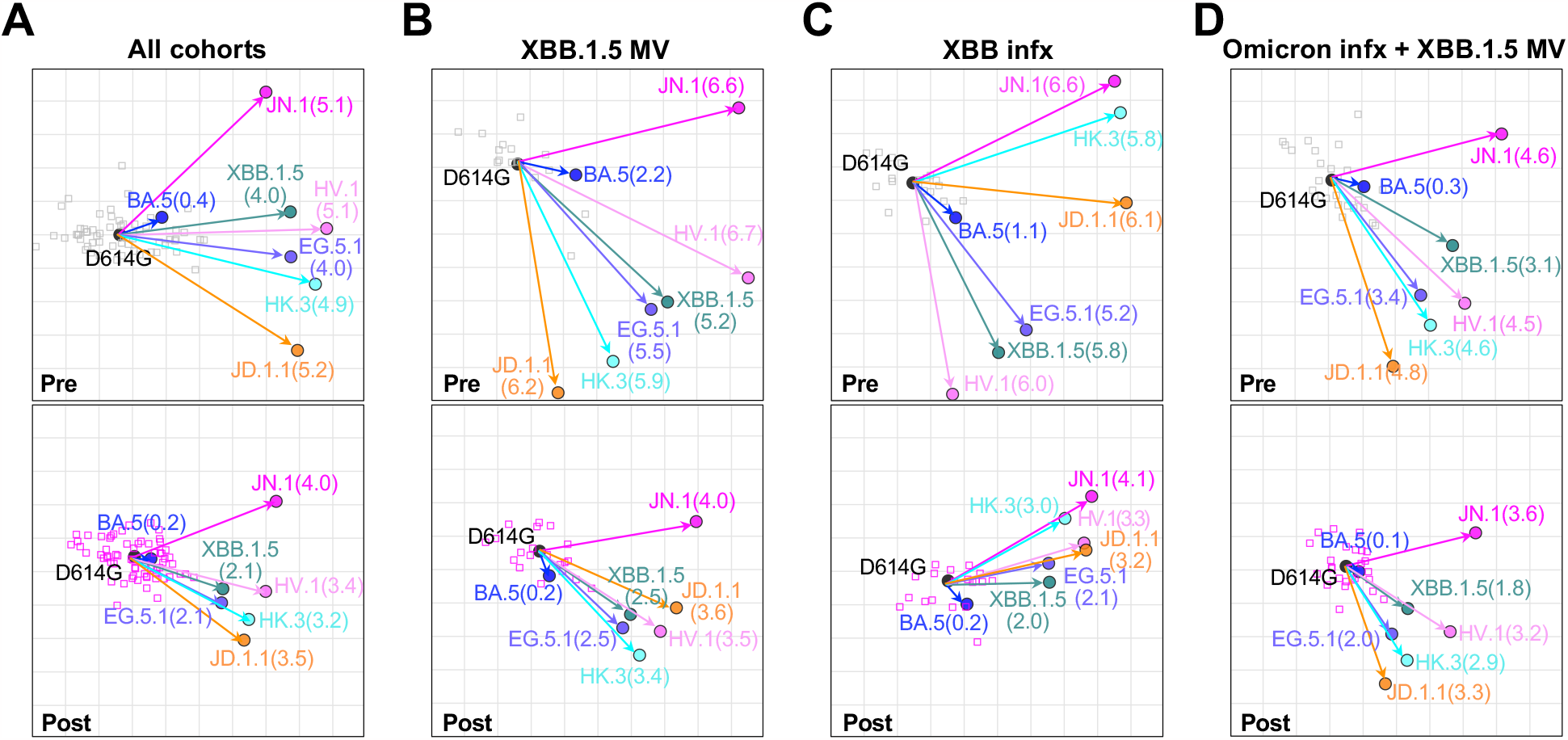
Antigenic cartography of serum virus-neutralizing data. Antigenic maps for all cohorts (**A**), the XBB.1.5 monovalent vaccine (XBB.1.5 MV) cohort (**B**), the XBB infection (XBB infx) cohort (**C**), and the infection + XBB.1.5 monovalent vaccine (Omicron infx + XBB.1.5 MV) cohort (**D**). The top row shows antigenic maps generated with pre-XBB sera, and the bottom row shows maps generated with post-XBB sera. The length of each square in the antigenic maps corresponds to one antigenic unit and represents an approximately 2-fold change in ID_50_ titer. Virus positions are shown in closed circles, while serum positions are shown by gray squares (pre-XBB sera) or pink squares (post-XBB sera). Antigenic distance from D614G is shown for each virus in parenthesis.

### Antigenic cartography

The serum neutralization data from all three cohorts combined, as well as individually, were used to construct antigenic maps (**Figures 3A-3D**), which graphically emphasize several key points. First, the discernible shortening of antigenic distances between D614G and other SARS-CoV-2 variants after a shot of XBB.1.5 monovalent vaccine (**Figures 3B and 3D**) was indicative of the significant boost in antibody potency and breadth. Second, the shortening of these antigenic distances after XBB.1.5 infection was also similar (**Figure 3C**) to that of XBB.1.5 vaccine booster (**Figure 3B**), suggesting that infection and vaccination resulted in comparable enhancement of antibody responses. Third, the emergent subvariants HV.1, HK.3, and JD.1.1 clustered together but were more distant than XBB.1.5 and EG.5.1 (**Figure 3**), demonstrating not only their antigenic similarity but also their greater antibody resistance compared to their predecessors. Lastly, JN.1 was antigenically distinct and more distant.

### Comparison of XBB.1.5 monovalent mRNA booster versus BA.5 bivalent mRNA booster

Following the XBB.1.5 monovalent vaccine booster, the highest neutralizing titers were observed against D614G and BA.5, not against XBB.1.5 (**Figures 2B and 2D**). This finding showed that there was considerable “back boosting” of antibodies directed to prior SARS-CoV-2 variants, which is likely the consequence of immunological imprinting (Koutsakos and Ellebedy, 2023) from prior vaccinations with the wildtype monovalent vaccine and the BA.5 bivalent vaccine. Nevertheless, an XBB.1.5 monovalent vaccine booster did markedly elevate serum neutralization titers against all Omicron subvariants tested (**Figures 2B and 2D)**, in contrast to prior results obtained after the BA.5 bivalent vaccine boost (Collier et al., 2023; Wang et al., 2023a; Wang et al., 2023b; Wang et al., 2023c; Wang et al., 2023e). We therefore compared the severity of immunological imprinting between XBB.1.5 monovalent vaccine and BA.5 bivalent vaccine. Serum neutralization data against D614G, BA.5, and XBB.1.5, generated using assays identical to those described herein, were extracted from our previous report (Wang et al., 2023a) on a cohort of individuals who received four shots of a wildtype monovalent vaccine followed by two shots of a BA.5 bivalent vaccine, and then compared with data extracted from two cohorts in the present study (**Figure 4A**). In individuals who received a second BA.5 bivalent booster, increases in mean serum neutralization titers against BA.5 were similar to that against D614G (2.6-fold versus 2.0-fold) (**Figure 4B**). However, strikingly, both the XBB.1.5 monovalent vaccine booster cohort (**Figure 4C**) and XBB breakthrough infection cohort (**Figure 4D**) showed markedly higher increases in mean neutralizing antibody titers against XBB.1.5 (27.0-fold and 28.6-fold, respectively) than against D614G (3.2-fold and 3.0-fold, respectively). These contrasting findings indicate that immunological imprinting is less severe for the XBB.1.5 monovalent vaccines.

**Figure 4.**
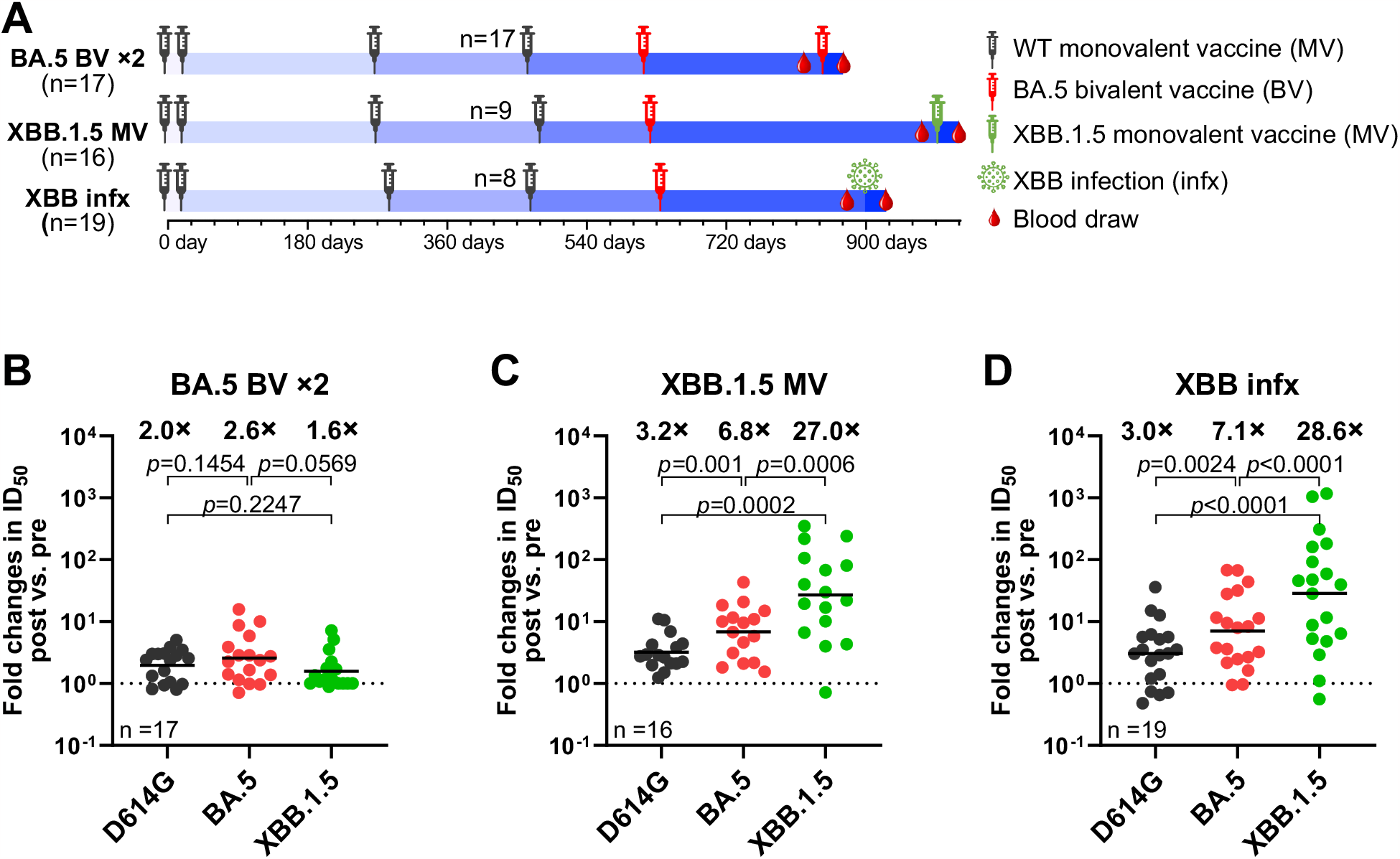
XBB.1.5 monovalent mRNA vaccines induced stronger boosts than a second BA.5 bivalent mRNA vaccine. **A**. Timeline representation of vaccine administration, SARS-CoV-2 infection, and serum collection intervals for each cohort. The cohort that received a second BA.5 bivalent vaccine (BA.5 BV x2) was previously described (Wang et al., 2023a). Indicated timepoints represent the median in days for each cohort, with day 0 defined as the day of the initial SARS-CoV-2 vaccination. Numbers of participants for each group receiving a fourth wildtype (WT) monovalent vaccine is indicated. n, sample size. **B-D**. Fold changes in ID_50_ titers of the indicated cohorts against D614G, BA.5, and XBB.1.5 between pre and post vaccination or infection. Geometric mean fold changes in ID_50_ titer are shown as black bars and denoted above the dots. Statistical analyses were performed by employing Wilcoxon matched-pairs signed-rank tests. Data for the BA.5 BV x2 cohort were extracted from a previously published study (Wang et al., 2023a).

## Discussion

Our findings showed that both XBB.1.5 monovalent mRNA vaccine booster and XBB.1.5 breakthrough infection markedly increased the magnitude of serum neutralizing antibodies against currently prevalent SARS-CoV-2 Omicron subvariants such as XBB.1.5 and EG.5.1 (**Figure 2**), in general agreement with clinical data posted by Chalkias et al (Chalkias et al., 2023), Stankov et al (Stankov et al., 2023), and Kosugi et al (Kosugi et al., 2023), and animal immunization results posted by Patel et al (Patel et al., 2023), and Modjarrad et al (Modjarrad et al., 2023). The latter three studies also found that there are strong specific T-cell responses directed to the spike protein of XBB subvariants (Modjarrad et al., 2023; Patel et al., 2023; Stankov et al., 2023). Here, we extended our study to include emerging Omicron subvariants that are now gaining traction and expanding rapidly, including HV.1, HK.3, JD.1.1, which are descendants of the XBB lineage, as well as JN.1, which is closely related to BA.2.86 (**Figure 1)**. Serum neutralizing titers against these emergent viruses increased by ∼13-to-27-fold after an XBB.15 monovalent vaccine booster in individuals without an infection history (**Figure 2B**), and by ∼10-fold in individuals with a prior Omicron infection (**Figure 2D**). Interestingly, we also showed that those boosted by an XBB.1.5 monovalent vaccine elicited serum neutralization potency and breadth similar to those with an XBB.1.5 breakthrough infection (**Figures 2B, 2C, 3B, and 3C**).

Our results also showed that HV.1, HK.3, and JD.1.1 are more resistant to serum neutralization than XBB.15 by about 1.9-to-2.8-fold (**Figures 2B-2D**), a finding that suggests that these emergent subvariants are likely to have a growth advantage in the population over their immediate precursors. If so, we can expect these new sublineages to replace XBB.1.5 and EG.5.1. Likewise, JN.1 is even more antibody resistant, by 2.9-to-4.3-fold, to the serum samples tested here (**Figure 3**). Widespread application of the updated XBB.1.5 monovalent vaccines could confer an even larger growth advantage in the population to JN.1 as well as to the related BA.2.86, thereby posing a potential threat to the newly authorized COVID-19 vaccines.

Our findings suggest that immunological imprinting is evident with the XBB.1.5 monovalent mRNA vaccines studied, in concordance with findings by Tortorici et al. (Tortorici et al., 2023). However, as discussed above, it is not nearly as severe as those observed for the BA.5 bivalent vaccines (**Figure 4**). One potential explanation is that XBB.1.5 is genetically and antigenically more distant from the ancestral SARS-CoV-2 than BA.5, which might mitigate immunological imprinting to an extent. Perhaps a more likely explanation is the non-inclusion of the ancestral spike in the current XBB.1.5 monovalent vaccines. Previous studies on the bivalent WA1+BA5 vaccines by our team (Wang et al., 2023a; Wang et al., 2023b; Wang et al., 2023c; Wang et al., 2023e) and others (Collier et al., 2023) suggested that the inclusion of the ancestral spike exacerbated the problem of imprinting and recommended its removal. Our findings herein indicate that WHO, FDA, and the vaccine manufacturers made the right choice by formulating the new COVID-19 vaccines based on XBB.1.5 spike alone, without including the ancestral spike.

This study is limited to evaluation of serum neutralizing antibodies, without addressing T-cell responses (Sette and Crotty, 2021; Vogel et al., 2021; Zhang et al., 2022) or mucosal immunity (Afkhami et al., 2022; Mao et al., 2022; Tang et al., 2022), both of which could provide added protection against SARS-CoV-2. Moreover, we have only examined acute antibody responses after XBB.1.5 monovalent vaccine booster or XBB.1.5 infection, but how such responses evolve over time will require follow-up studies. These limitations notwithstanding, our results not only demonstrate that administration of an XBB.1.5 monovalent mRNA vaccine booster can elicit robust neutralizing antibodies against current and emerging SARS-CoV-2 variants, but also support FDA’s recommendation to apply these updated COVID-19 vaccines more widely to confer greater protection to the public.

## STAR METHODS

Detailed methods are provided in the online version of this paper and include the following:

- **KEY RESOURCES TABLE**
- **RESOURCE AVAILABILITY**
  - Lead contact
  - Materials availability
  - Data and code availability
- **EXPERIMENTAL MODEL AND SUBJECT DETAILS**
  - Clinical cohorts
  - Cell lines
- **METHOD DETAILS**
  - Pseudovirus neutralization assay
  - Phylogenetic analysis
  - Antigenic cartography
- **QUANTIFICATION AND STATISTICAL ANAYLYSIS**

## ACKNOWLEDGEMENTS

This study was supported by funding from the NIH SARS-CoV-2 Assessment of Viral Evolution (SAVE) Program (Subcontract No. 0258-A709-4609 under Federal Contract No. 75N93021C00014) to D.D.H., as well as funding from the NIH contract 75N93019C00051 to A.G. We express our gratitude to Zijin Chu, Theresa Knowalski-Dobson, Emily Stoneman, David Manthei, Anna Buswinka, Gabe Simjanovski, Joseph Wendzinski, Mayurika Patel, Kathleen Lindsey, Dawson Davis, Victoria Blanc, Savanna Sneeringer, and Pamela Bennett-Baker of the IASO study team for conducting the IASO study.

## AUTHOR CONTRIBUTIONS

The study was conceptualized by A.G., L.L. and D.D.H. Experiments were conducted and data analyzed by Q.W., L.L., Y.G., I.A.M., and A.B. Project management was handled by Q.W. Serum samples were collected by R.V., C.G., A.G., and their colleagues. The results were analyzed and the manuscript was written by Q.W., Y.G., L.L., and D.D.H. All contributing authors have reviewed and endorsed the manuscript.

## Figures and legends

**Table S1.**
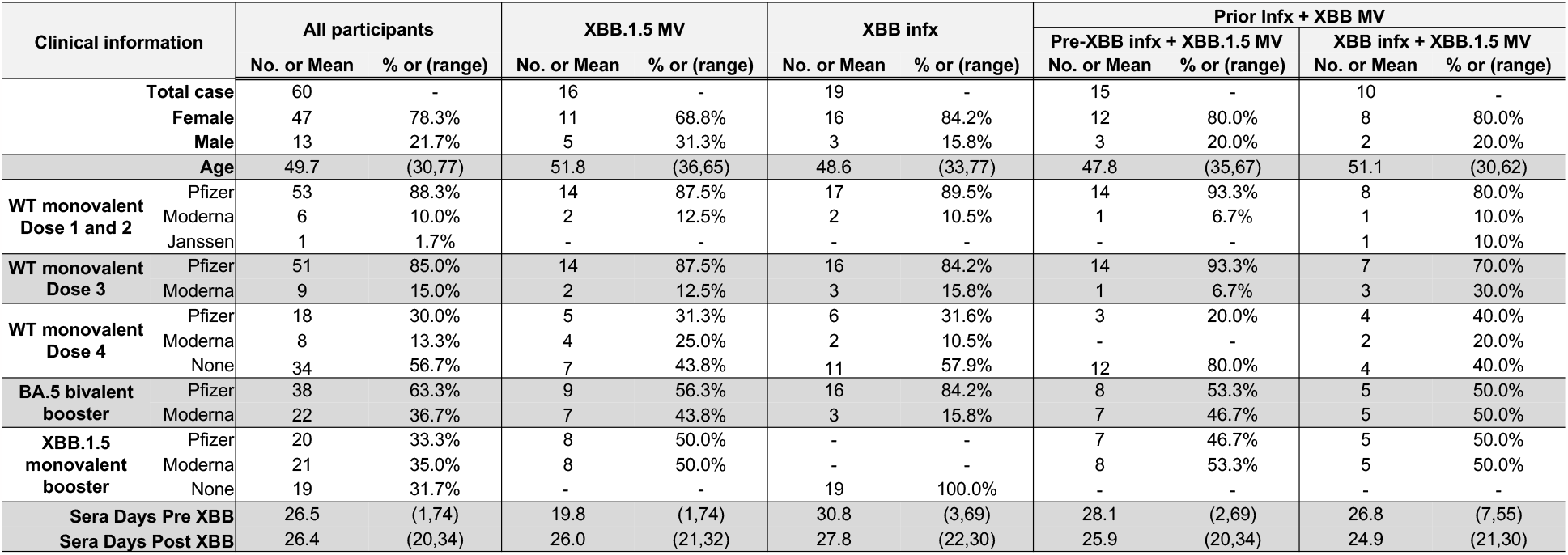
Summarized participant information. Demographics, vaccines, and serum collection information are summarized for each cohort. Listed values represent the mean and range (age and sera collection variables) or number and percentage (vaccine type and sex variables). See also **Figure 2**.

| Clinical information |  | All participants |  | XBB.1.5 MV |  | XBB infx |  | Prior Infx + XBB MV |  |  |  |
| --- | --- | --- | --- | --- | --- | --- | --- | --- | --- | --- | --- |
|  |  | No. or Mean | % or (range) | No. or Mean | % or (range) | No. or Mean | % or (range) | Pre-XBB infx + XBB.1.5 MV |  | XBB infx + XBB.1.5 MV |  |
| Total case |  | 60 | - | 16 | - | 19 | - | 15 | - | 10 | - |
| Female |  | 47 | 78.3% | 11 | 68.8% | 16 | 84.2% | 12 | 80.0% | 8 | 80.0% |
| Male |  | 13 | 21.7% | 5 | 31.3% | 3 | 15.8% | 3 | 20.0% | 2 | 20.0% |
| Age |  | 49.7 | (30,77) | 51.8 | (36,65) | 48.6 | (33,77) | 47.8 | (35,67) | 51.1 | (30,62) |
| WT monovalent<br>Dose 1 and 2 | Pfizer | 53 | 88.3% | 14 | 87.5% | 17 | 89.5% | 14 | 93.3% | 8 | 80.0% |
|  | Moderna | 6 | 10.0% | 2 | 12.5% | 2 | 10.5% | 1 | 6.7% | 1 | 10.0% |
|  | Janssen | 1 | 1.7% | - | - | - | - | - | - | 1 | 10.0% |
| WT monovalent<br>Dose 3 | Pfizer | 51 | 85.0% | 14 | 87.5% | 16 | 84.2% | 14 | 93.3% | 7 | 70.0% |
|  | Moderna | 9 | 15.0% | 2 | 12.5% | 3 | 15.8% | 1 | 6.7% | 3 | 30.0% |
| WT monovalent<br>Dose 4 | Pfizer | 18 | 30.0% | 5 | 31.3% | 6 | 31.6% | 3 | 20.0% | 4 | 40.0% |
|  | Moderna | 8 | 13.3% | 4 | 25.0% | 2 | 10.5% | - | - | 2 | 20.0% |
|  | None | 34 | 56.7% | 7 | 43.8% | 11 | 57.9% | 12 | 80.0% | 4 | 40.0% |
| BA.5 bivalent<br>booster | Pfizer | 38 | 63.3% | 9 | 56.3% | 16 | 84.2% | 8 | 53.3% | 5 | 50.0% |
|  | Moderna | 22 | 36.7% | 7 | 43.8% | 3 | 15.8% | 7 | 46.7% | 5 | 50.0% |
| XBB.1.5<br>monovalent<br>booster | Pfizer | 20 | 33.3% | 8 | 50.0% | - | - | 7 | 46.7% | 5 | 50.0% |
|  | Moderna | 21 | 35.0% | 8 | 50.0% | - | - | 8 | 53.3% | 5 | 50.0% |
|  | None | 19 | 31.7% | - | - | 19 | 100.0% | - | - | - | - |
| Sera Days Pre XBB |  | 26.5 | (1,74) | 19.8 | (1,74) | 30.8 | (3,69) | 28.1 | (2,69) | 26.8 | (7,55) |
| Sera Days Post XBB |  | 26.4 | (20,34) | 26.0 | (21,32) | 27.8 | (22,30) | 25.9 | (20,34) | 24.9 | (21,30) |

**Table S2.**
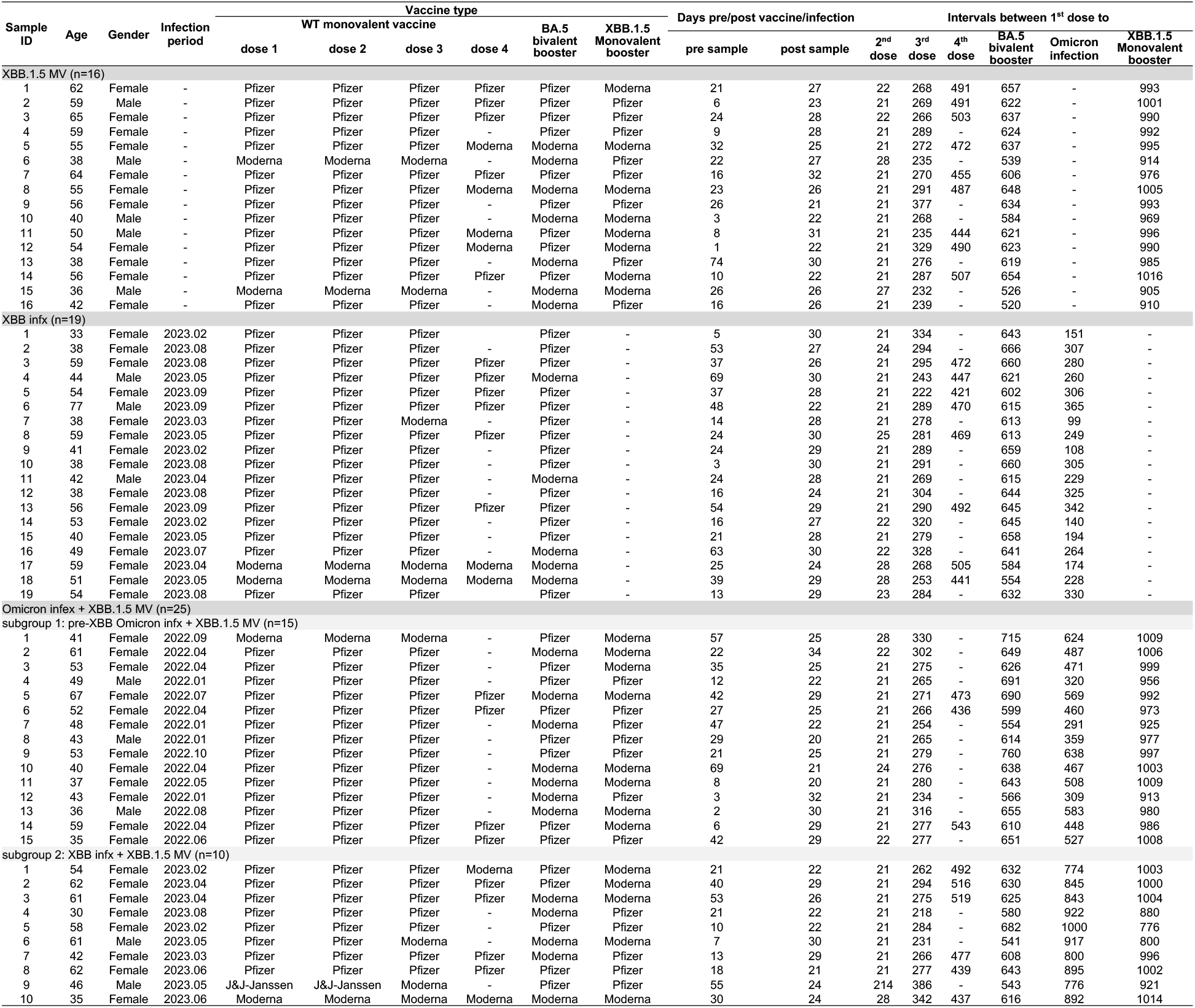
Participant details. Details are listed for each participant including demographics as well as vaccine, infection, and serum collection information. See also **Figure 2**.

**Figure S1.**
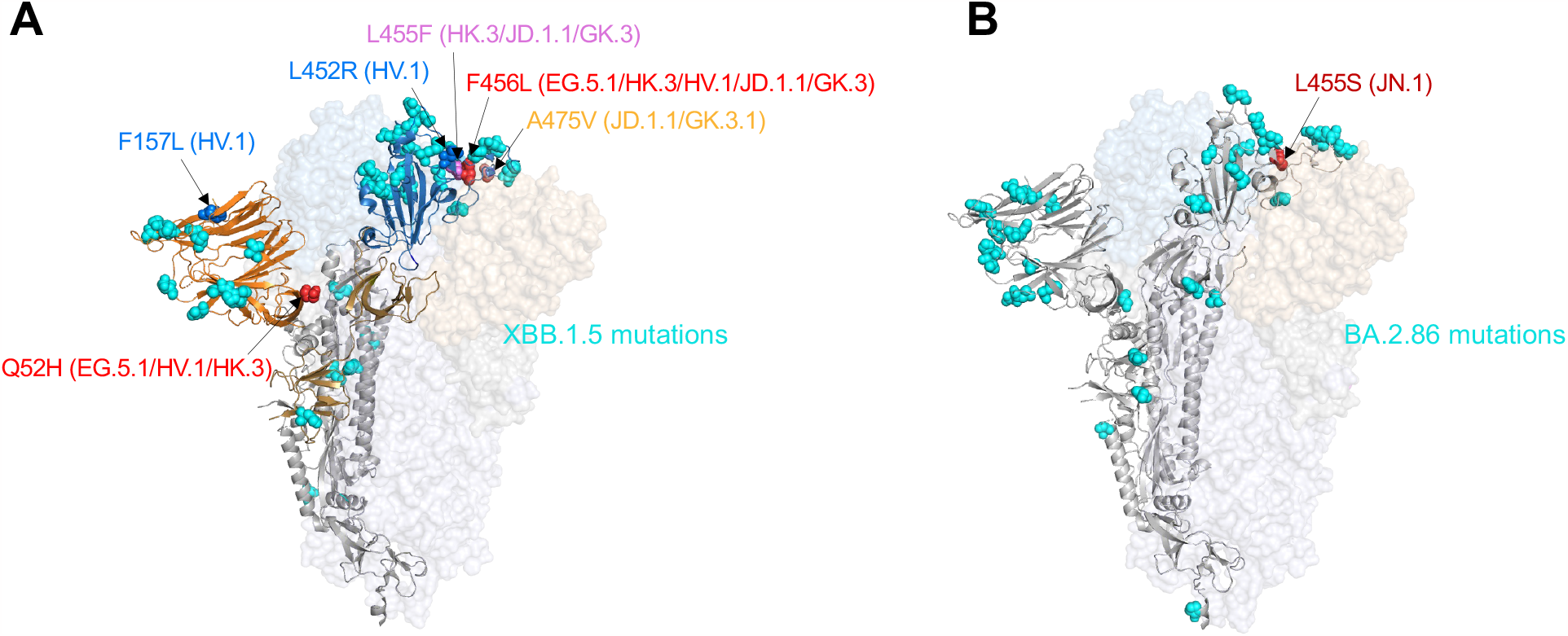
Spike mutations found in emerging SARS-CoV-2 Omicron subvariants. **A**. Mutations found in EG.5.1, HV.1, HK.3, JD.1.1, and GK.3 on top of the XBB.1.5 spike. **B**. Location of the L455S mutation in JN.1 on top of the BA.2.86 spike. Mutations present in XBB.1.5 and BA.2.86 are highlighted in cyan. The spike protein structure is obtained under PDB ID: 6ZGE (Wrobel et al., 2020). See also **Figure 1**.

**Figure S2.**
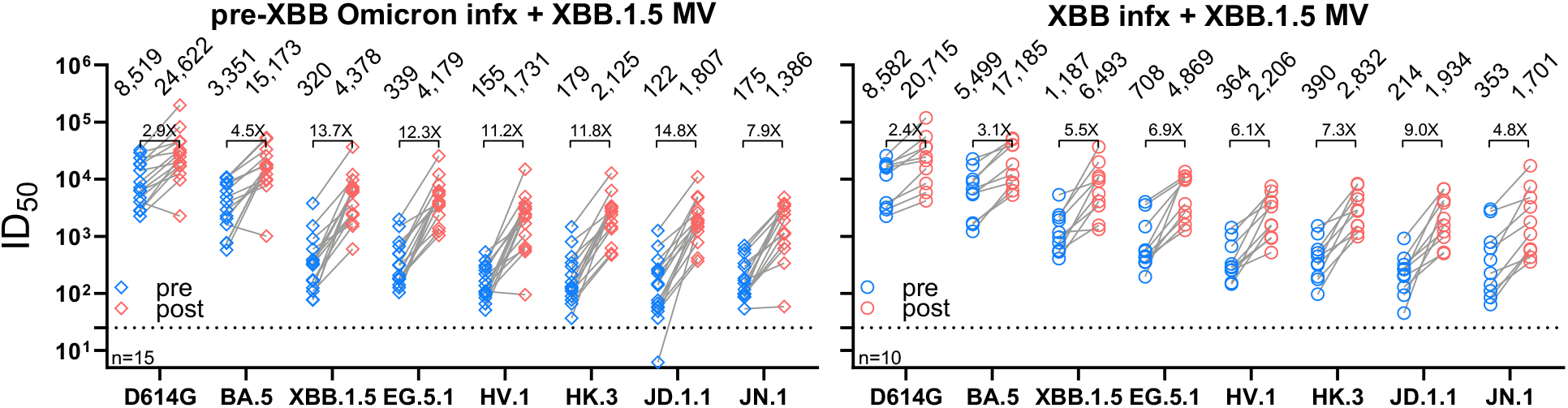
Neutralizing antibody titers before and after an XBB.1.5 mRNA vaccine booster following a pre-XBB Omicron infection or an XBB infection. Participants from the “Omicron infx + XBB.1.5 MV” cohort were stratified into two groups based on the infection strain. Geometric mean ID_50_ titers are shown along with the fold change between pre and post XBB.1.5 vaccination against each indicated virus. The dotted line represents the assay limit of detection of 25. “n” denotes the sample size. See also **Figure 2**.

**Figure S3.**
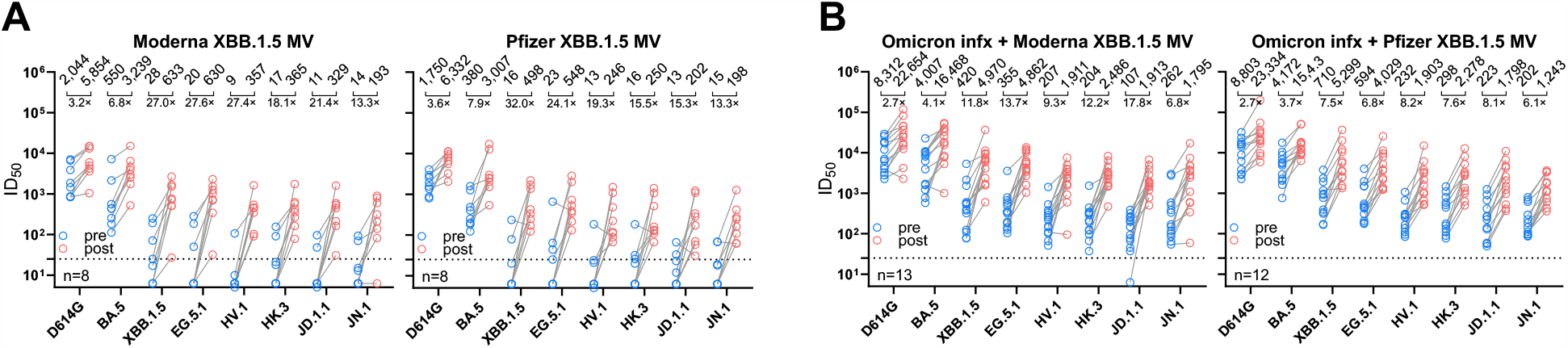
Neutralizing antibody titers before and after a Moderna or Pfizer XBB.1.5 mRNA vaccine booster. Participants from the “XBB.1.5 MV” cohort (**A**) and “Omicron infx + XBB.1.5 MV” (**B**) were stratified into two groups based on the vaccine manufacturer. Geometric mean ID_50_ titers are shown along with the fold change between pre and post XBB.1.5 vaccination against each indicated virus. The dotted line represents the assay limit of detection of 25. “n” denotes the sample size. See also **Figure 2**.

## RESOURCE AVAILABILITY

### Lead contact

Further information and requests for resources and reagents should be directed to and will be fulfilled by the Lead Contact Author David D. Ho.

### Materials availability

All reagents generated in this study are available from the Lead Contact with a completed Materials Transfer Agreement.

### Data and code availability

Data reported in this paper will be shared by the lead contact upon request. This paper does not report original code.

Any additional information required to reanalyze the data reported in this paper is available from the lead contact upon request.

## EXPERIMENTAL MODEL AND SUBJECT DETAILS

### Clinical cohorts

Longitudinal sera were obtained as part of a continuing cohort study, Immunity-Associated with SARS-CoV-2 Study (IASO), which began in 2020 at the University of Michigan in Ann Arbor, Michigan (Simon et al., 2022). Written informed consent was provided by all participants and sera were collected according to the protocol approved by the Institutional Review Board of the University of Michigan Medical School. Participants in the IASO study completed weekly symptom surveys and were tested for SARS-CoV-2 with any report of symptoms. All serum samples were examined by anti-nucleoprotein (NP) ELISA to confirm status of prior SARS-CoV-2 infection.

For this study, we included sera from 60 individuals in three distinct clinical cohorts: 1) individuals with no recorded SARS-CoV-2 infections who had received an XBB.1.5 monovalent vaccine booster (“XBB.1.5 MV”); 2) individuals with a recent XBB SARS-CoV-2 infection who had not received the XBB.1.5 booster (“XBB infx”); and 3) individuals with prior infection who also received the XBB.1.5 booster (“Omicron infx + XBB.1.5 MV”). The final cohort was divided into subgroup 1, with documented infection prior to 2023, and subgroup 2, with documented infection after February 2023. Individuals in all cohorts received either three or four doses of a wildtype monovalent vaccine as well as a single BA.5 bivalent booster.

Most participants were female (78.3%) with an average age of 49.7 years. Sera were collected an average of 26 days pre and post XBB.1.5 vaccination or XBB infection. Sera were examined by anti-nucleoprotein (NP) ELISA to determine status of prior SARS-CoV-2 infection. Demographic, vaccination, and serum collection details are summarized for each cohort and subgroup in **Table S1**, and details are shown for each participant in **Table S2**.

### Cell lines

293T (CRL-3216) and Vero-E6 (CRL-1586) cells were obtained from ATCC and cultured in the conditions following manufacturer’s instructions. The morphology of each cell line was visually confirmed before use. All cell lines tested negative for mycoplasma.

## METHOD DETAILS

### Pseudovirus neutralization assay

Plasmids encoding SARS-CoV-2 variant spikes, including D614G, BA.5, XBB.1.5, and EG.5.1, were generated in previous studies (Wang et al., 2023b; Wang et al., 2022; Wang et al., 2023f; Wang et al., 2023g). Plasmids expressing HV.1, HK.3, JD.1.1, and JN.1 spikes were generated by introducing mutations to the XBB.1.5 (Wang et al., 2023b), EG.5.1 (Wang et al., 2023f) or BA.2.86 (Wang et al., 2023d) spike (**Figure 1C**) using the QuikChange® mutagenesis kit.

To produce pseudotyped viruses of SARS-CoV-2 variants, 293T cells were transfected with the spike-encoding plasmids described above using 1 mg/mL PEI (Polyethylenimine). One day post-transfection, the 293T cells were then incubated with VSVG*ΔG-luciferase (Kerafast, Inc.) at a multiplicity of approximately 3 to 5 for 2 hours followed by three washes with PBS. The cells were then cultured with fresh medium for an additional day. Cell supernatants containing viruses were collected, clarified by centrifugation, aliquoted, and stored at -80°C until use.

The viral titer of each variant was titrated and normalized for the neutralization assays. Serum samples were diluted in triplicate in 96-well plates, starting from a 12.5-fold dilution, and then incubated with an equal volume of virus for 1 hour at 37°C before adding 2 × 10^4^ cells/well of Vero-E6 cells. The cells were then cultured overnight, harvested, and lysed for measurement of luciferase activity using SoftMax Pro v.7.0.2 (Molecular Devices). Reductions in luciferase activity at given dilutions of sera were calculated, and ID_50_ values of sera were obtained by fitting the virus-reduction data using a non-linear five-parameter dose-response curve in GraphPad Prism V.10.

### Phylogenetic analysis

Genome sequences of SARS-CoV-2 subvariants are retrieved from the GISAID database (Elbe and Buckland-Merrett, 2017). The spike protein sequences are then extracted from these genomes using an in-house Python script. Post-extraction, these sequences are aligned by MUSCLE software, version 3.8.31. Sequencing sites with low quality, identified by the presence of ‘N’, underwent a manual curation to align the mutations with the consensus for each variant. A Maximum-Likelihood phylogenetic tree was constructed with MEGA11 software, utilizing the Tamura-Nei model, and its robustness was verified through 500 bootstrap replications.

### Antigenic cartography

The antigenic distances between serum samples and D614G, along with other SARS-CoV-2 variants, were calculated by integrating the ID_50_ values of individual serum samples using a published antigenic cartography method (Smith et al., 2004). Visualizations are created with the Racmacs package (version 1.1.4, https://acorg.github.io/Racmacs/) within R software version 4.0.3. The optimization is set to 2,000 steps, with the “minimum column basis” parameter set to “none”. The “mapDistances” function was used to calculate the antigenic distances, with the average distances from all serum samples to each variant representing the final outputs. For each group, D614G was positioned as the center point of the sera. The seeds for each antigenic map are manually adjusted to position D614G left horizontally in relation to other variants.

## QUANTIFICATION AND STATISTICAL ANAYLYSIS

Serum neutralization ID_50_ values were calculated using a five-parameter dose-response curve in GraphPad Prism v.10. Evaluations of statistical significance were performed employing two-tailed Wilcoxon matched-pairs signed-rank tests using GraphPad Prism v.10 software.

